# Actinomarina, resolved

**DOI:** 10.64898/2026.04.03.716425

**Authors:** Torben N. Nielsen, Lauren M. Lui

**Author notes:** Both authors contributed equally.

## Abstract

*Actinomarina* (*Ca.* Actinomarina minuta) is among the smallest known free-living bacteria and among the most abundant photoheterotrophs in ocean surface waters, yet not a single complete genome has been published for any member of the order Actinomarinales. Here we present 84 complete *Actinomarina* genomes assembled from Oxford Nanopore metagenomes of the San Francisco Estuary (SFE), together with 29 additional high-quality (≥95% complete, <5% contamination, single contig) non-circular assemblies. These genomes define 9 species at 95% ANI, three of which are novel, and represent, to our knowledge, the first complete genomes for the entire Actinomarinales order. An expanded phylogeny incorporating 23 high-quality NCBI genomes and a Casp-actino5 outgroup confirms species boundaries and places NCBI genomes among SFE clades.

The pangenome of 9,278 clusters is approaching closure (decay ratio 0.55), with a core genome of 227 single-copy genes (2.4%). Singletons (3,858 clusters, 42%) are concentrated in a tRNA-bounded hypervariable region (HVR) at 85–90% of the chromosome from dnaA — a similar replicative distance to the HVR recently described in *Pelagibacter* (7–15% from dnaA; Lui & Nielsen, in preparation) but on the opposite replichore. The HVR is flanked by tRNA genes at both boundaries, and in 32 of 84 genomes, the selenocysteine tRNA (selC) is physically located inside the HVR.

All 84 genomes encode selenocysteine tRNA (selC) and the selenocysteine biosynthesis genes selA and selD; the elongation factor selB is present but divergent. This has not been previously reported for *Actinomarina*. Deep learning selenoprotein prediction (deep-Sep) identifies ∼5 selenoproteins per genome in 69 families, including a selenoprotein form of selD itself. Retention of dedicated Sec biosynthetic machinery serving multiple targets in a genome of ∼1.1 Mbp implies that the catalytic advantage of selenocysteine over cysteine is large enough to justify the cost.

KofamScan pathway reconstruction reveals the most extensive set of auxotrophies yet documented from complete genomes in a free-living marine bacterium: arginine, histidine, tryptophan, and thiamine biosynthesis are universally absent, biotin biosynthesis is non-functional (final step absent), and only 5 of 8 TCA cycle steps are retained at the standard detection threshold. Gene order is extensively rearranged between species: no gene adjacency is universal across all 84 genomes. NCBI currently lists 396 *Actinomarina* genomes, none complete; of the 39 that pass quality thresholds, 41% are misclassified by NCBI, belonging to other genera by GTDB-Tk reclassification.

## Introduction

*Candidatus* Actinomarina minuta is among the smallest known free-living bacteria, with cell volumes of approximately 0.013 µm³ (Ghai et al., 2013). It is the dominant member of the order Actinomarinales (phylum Actinomycetota, class Acidimicrobiia), a lineage that constitutes up to 4% of prokaryotic cells in the ocean’s photic zone — second only to SAR11 among heterotrophic and photoheterotrophic bacteria at the deep chlorophyll maximum (Ghai et al., 2013; Lopez-Perez et al., 2020). *Actinomarina* is a photoheterotroph, using a type I actinorhodopsin for light-driven proton pumping (Lopez-Perez et al., 2020) and encoding a heliorhodopsin of unknown function (Pushkarev et al., 2018). Its genomes are among the smallest known for any free-living organism: approximately 1.1 Mbp, with GC content around 33%, coding densities of approximately 97%, and a median intergenic spacer of just 3 bp.

Despite this ecological prominence, the genomic diversity of *Actinomarina* remains poorly resolved. Lopez-Perez et al. (2020) characterized 182 genomes from single-amplified genomes (SAGs) and metagenome-assembled genomes (MAGs) across the Actinomarinales, defining 5 genera and 18 genomospecies. However, none of these genomes were complete. NCBI currently lists 396 genomes classified as *Actinomarina*, all contig-level MAGs, with no complete (circular) assembly for any member of the order. Without complete genomes, every apparent gene absence is ambiguous: a missing pathway step may reflect genuine biological gene loss, or it may be an assembly gap, a binning error, or incomplete SAG amplification. For a genome of only ∼1.1 Mbp, where a single missing gene can change the metabolic interpretation, this ambiguity is especially consequential.

The difficulty of assembling complete *Actinomarina* genomes from metagenomes has two likely causes. First, multiple *Actinomarina* species co-occur in the same sample with high core gene similarity, creating regions of near-identity that collapse in assembly graphs. Second, the hypervariable region — first noted by Lopez-Perez et al. (2020) and characterized in detail here — carries genome-specific content that creates branching paths at a locus flanked by conserved sequence. Together, these create regions where conserved sequence flanks genome-specific content, which assemblers cannot resolve without sufficient coverage. Additionally, the extensive gene order rearrangement between species (described below) means that even outside the HVR, co-occurring species share conserved genes in different genomic contexts, multiplying the number of branching points in the assembly graph. Internal repeats are not a significant factor for long-read assembly: the longest repeat is 777 bp (median 152 bp), well within the span of individual ONT reads.

This study is the first in a series of companion papers characterizing the 20 most abundant bacterial genera in the SFE, each from complete genomes (biogeography paper in preparation). Here we exploit deep Oxford Nanopore metagenomes of the San Francisco Estuary, assembled with myloasm (Shaw, Marin & Li, 2026), to recover 84 complete *Actinomarina* genomes — the first complete genomes for any member of the Actinomarinales. Together with 29 additional high-quality non-circular assemblies and 23 confirmed *Actinomarina* genomes from NCBI, this collection allows us to address three questions. First, what is the species-level diversity of *Actinomarina* in the SFE, and how does it compare to the SAG-based descriptions of Lopez-Perez et al. (2020)? Second, what is the definitive metabolic capability of this lineage — which auxotrophies are real, and which were artifacts of incomplete assemblies? Third, does *Actinomarina* share the hypervariable region architecture recently described in *Pelagibacter*, and if so, what does this imply about convergent genome organization in small-genome marine bacteria?

## Results

### Recovery of 84 complete *Actinomarina* genomes

We assembled 84 complete (circular) *Actinomarina* genomes from ONT metagenomes of the SFE (Supplementary Table 1; Figure 1A). An additional 29 genomes met our high-quality thresholds (single-contig, ≥90% completeness, <5% contamination by CheckM2) but were not circular; these were included in phylogenomic analyses but excluded from pangenome, synteny, and chromosomal position analyses that require unambiguous gene order. All 113 genomes were classified as *Actinomarina* by GTDB-Tk (GTDB R226).

**Figure 1A:**
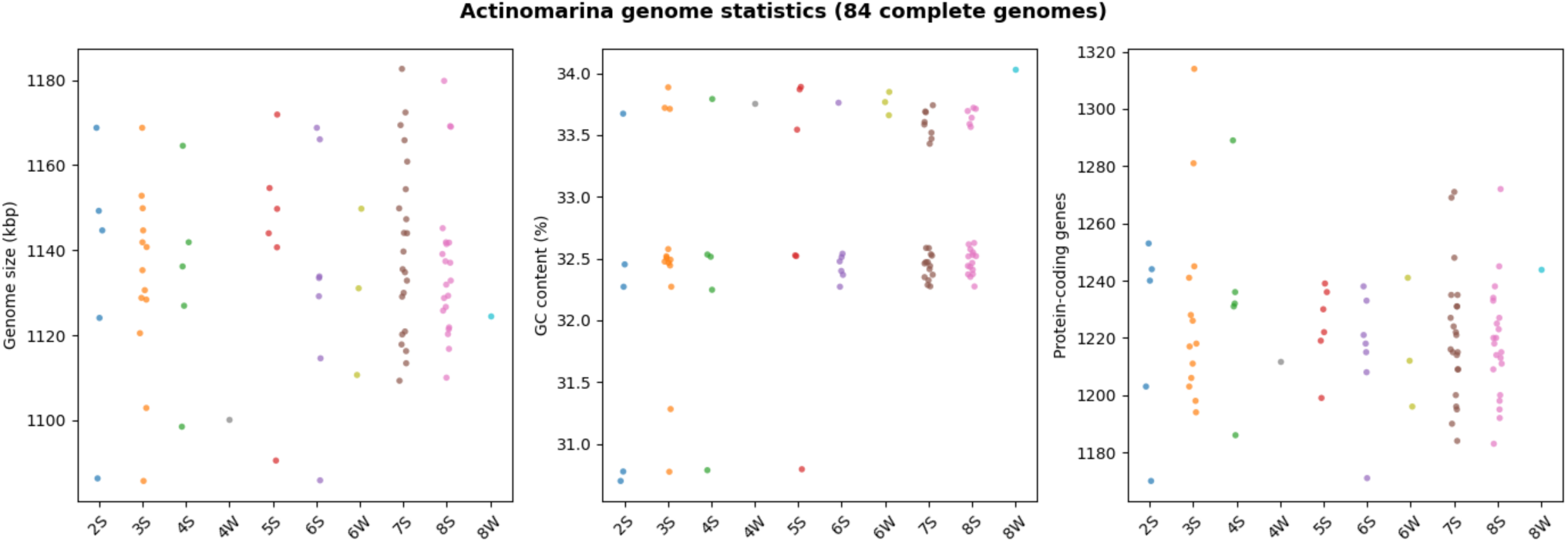
Distribution of genome sizes, GC content, and gene counts across the 84 complete genomes, colored by station.

The 84 complete genomes ranged in size from 1,090,312 to 1,184,021 bp (mean 1,130,456 bp), with GC content of 31.0–34.0% (mean 32.8%). Pyrodigal v3.6.3 (Larralde, 2022) in single-genome mode predicted 1,150 to 1,284 protein-coding genes per genome (mean 1,222), with coding densities of 97.1 ± 0.3%. Genomes were reoriented to begin at dnaA using dnaapler v1.3.0 (Bouras et al., 2024). Throughout this study, taxonomic assignments follow GTDB (Parks et al., 2025) via GTDB-Tk, which provides rank-normalized, phylogenetically consistent classifications; NCBI taxonomy does not distinguish genera within the Actinomarinales. Of the 29 non-circular genomes, 27 were successfully reoriented; 2 (2S_u7034620, 8S_u23307925) lacked dnaA entirely, consistent with a missing ∼10% region that includes the replication origin — likely an assembly tangle at a conserved locus shared with co-occurring species.

The 84 complete genomes were recovered from 10 of 16 SFE samples: 7S (22), 8S (21), 3S (13), 6S (7), 5S (6), 2S (5), 4S (5), 6W (3), 4W (1), and 8W (1). Summer stations contributed 79 of 84 genomes. No complete genomes were recovered from station 1 (freshwater) or from 5W, 7W, 3W, or 2W, consistent with *Actinomarina* being restricted to the mesohaline and polyhaline portions of the estuary.

tRNA gene prediction with tRNAscan-SE identified 39–52 tRNAs per genome (mean 44). All 84 genomes encode tRNAs for all 20 standard amino acids. The most redundant tRNAs are Leu (4.3 copies/genome), Arg (4.1), Ser (3.7), and Val (3.7). All 84 genomes also encode a selenocysteine tRNA (selC), a finding we discuss in detail below.

These 84 genomes represent the first complete assemblies for any member of the Actinomarinales. No complete genome for *Actinomarina* or any related genus existed in NCBI or any other public database as of March 2026.

### Nine species, three of which are novel

Pairwise ANI values were computed for all 113 genomes using skani. At the standard 95% ANI threshold, the 113 genomes clustered into 9 species-level groups (Figure 1B). Cluster sizes were 28, 24, 18, 12, 12, 9, 7, 2, and 1 — a markedly uneven distribution dominated by three large clusters that together account for 70 of 113 genomes (62%). Only one species is represented by a single genome.

**Figure 1B:**
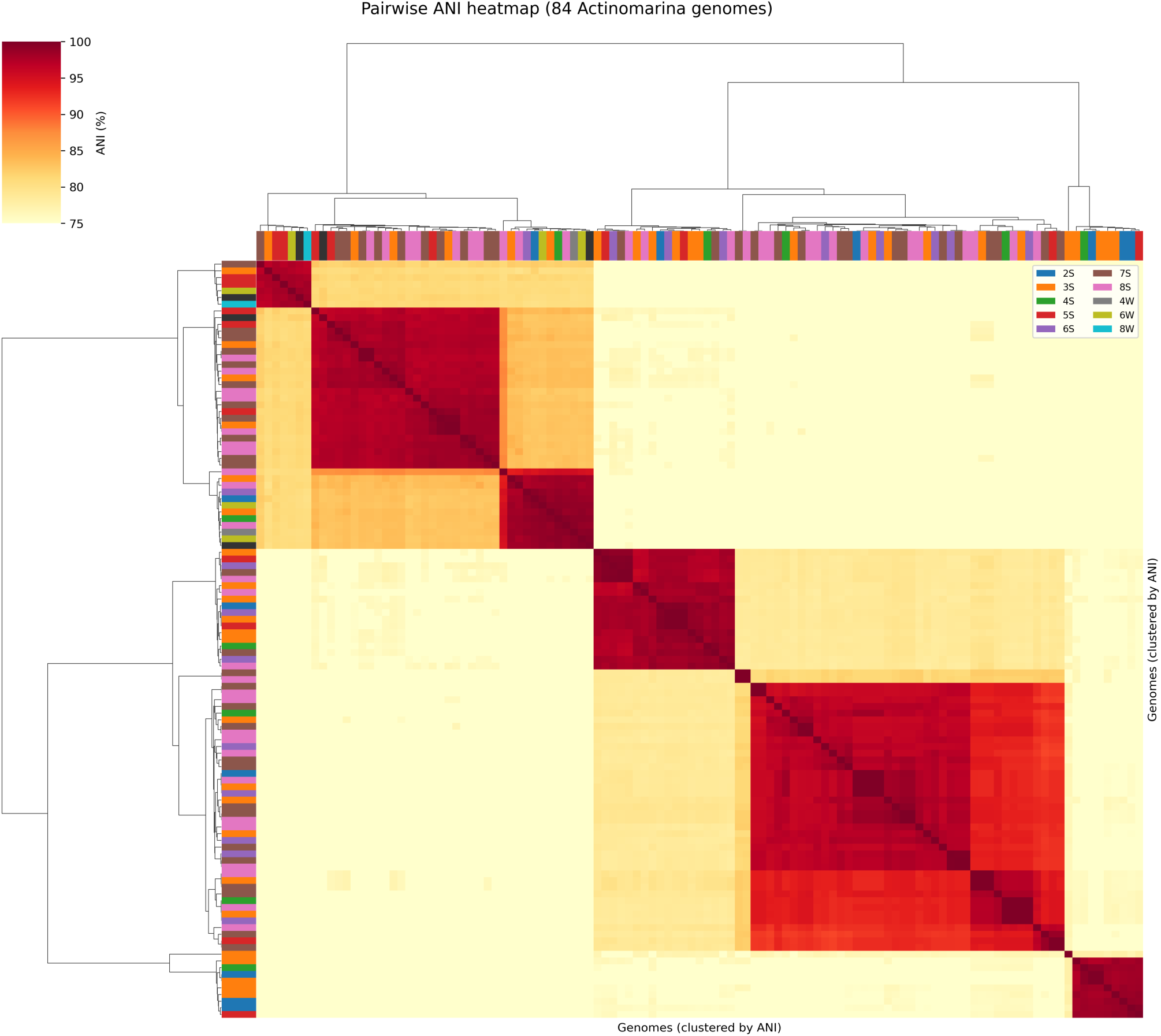
Pairwise ANI heatmap of 84 Actinomarina genomes ordered by hierarchical clustering.

GTDB-Tk classification assigned 6 of the 9 species to named GTDB species; the remaining 3 are novel (no match to any existing GTDB species). Every species has at least one complete (circular) genome, and the three largest clusters have 23, 15, and 16 complete genomes respectively. This pattern — few species, each deeply sampled — yields a mean of 12.6 genomes per species across all 113 genomes (9.3 per species among the 84 complete genomes), enabling robust within-species comparisons.

### NCBI genome comparison

NCBI currently lists 396 genomes classified as *Actinomarina*, all contig-level MAGs with no complete assemblies. CheckM2 assessment revealed that only 39 (10%) meet high-quality standards. GTDB-Tk reclassification of these 39 showed that 16 (41%) are misclassified — belonging to genera including *Nocardioides* (Propionibacteriales), *Casp-actino5* and *UBA11606* (Acidimicrobiales), *Alteriqipengyuania* (Sphingomonadales), and *Luminiphilus* (Pseudomonadales), with genome sizes of 1.6–4.0 Mbp that are incompatible with the ∼1.1 Mbp *Actinomarina* genome. Only 23 of 396 NCBI genomes both meet CheckM2 quality thresholds and are genuinely *Actinomarina* (Supplementary Table 2). Of these, 11 fall within our 9 species at ≥95% ANI, 5 represent different species at 80–95% ANI, and 7 are below the ANI detection threshold. Our 84 complete genomes outnumber the confirmed high-quality MAGs at NCBI by nearly 4:1.

### Phylogenomic structure

Phylogenomic inference was performed on a supermatrix of 227 single-copy core protein clusters (present in exactly one copy in all 84 complete genomes), aligned with MAFFT and concatenated into 65,482 amino acid positions. IQ-TREE analysis with per-gene LG+F+G4 partitions and 1000 ultrafast bootstrap replicates produced a well-resolved 84-taxon tree: 69% of internal branches (56/81) received ≥95% UFBoot support, 89% received ≥80%, and only 2 fell below 50%.

An expanded tree was inferred from 136 taxa (84 circular + 29 non-circular + 23 NCBI), using the same core genes with new sequences added via MAFFT –add –keeplength. Three non-circular genomes (3S_u19197949, 3S_u20094592, 8S_u23307925) had 11–14% gaps in the supermatrix due to missing core genes; these were retained with all-gap positions for missing genes. The rooted tree shows comparable support: 70% of internal branches (94/134) ≥95% UFBoot, 86% ≥80%, and 2 below 50%.

The tree was rooted using 3S_u11376018, a member of Casp-actino5 (order Acidimicrobiales, the sister order to Actinomarinales in GTDB). This outgroup retains 164 of the 227 core genes at ≥30% amino acid identity and 50% coverage (median identity 40.7%), with 28% total gaps in the supermatrix — distant but alignable. The rooted 137-taxon tree (Figure 2) confirms the monophyly of all 9 species clusters. NCBI genomes — from the Mediterranean, Tara Oceans, and the North Sea — are interspersed among SFE genomes rather than forming separate clades, with only one minor clade lacking SFE representation. This pattern indicates substantial overlap between SFE and globally sampled *Actinomarina* diversity, though geographic coverage of available NCBI genomes remains limited.

**Figure 2:**
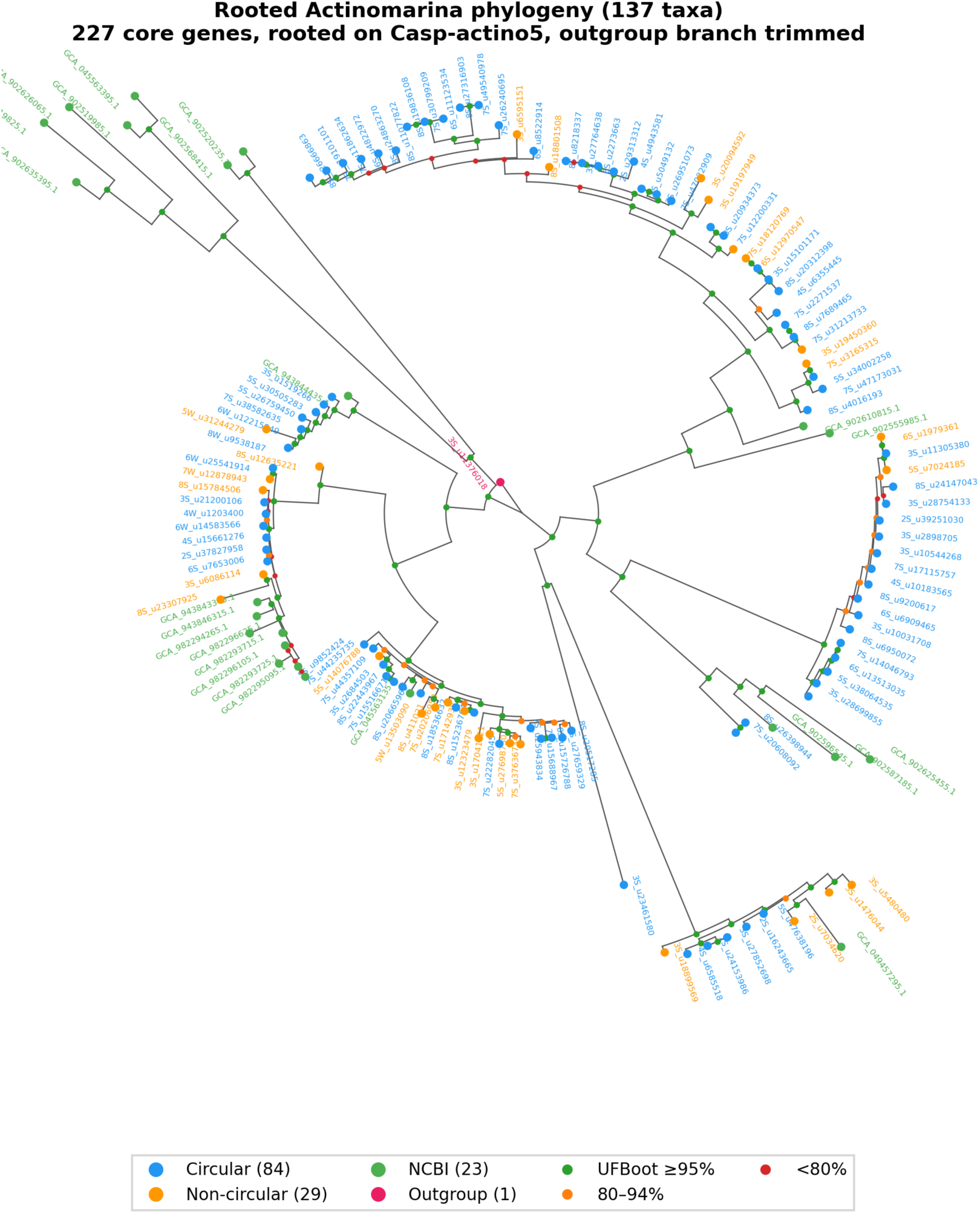
Rooted phylogenomic tree of 136 *Actinomarina* genomes plus Casp-actino5 outgroup (84 circular + 29 non-circular + 23 NCBI). Bootstrap support shown as colored dots. Tips colored by source. Outgroup branch trimmed for display.

### Pangenome architecture

MMseqs2 clustering of 102,662 predicted proteins from all 84 complete genomes at 70% identity and 80% bidirectional coverage yielded 9,278 gene clusters. Of these, 227 (2.4%) were strictly core — present in all 84 genomes (Supplementary Table 3) — and all 227 occur in a single copy per genome. The shell (present in 2 to 83 genomes) comprised 5,193 clusters, and 3,858 clusters (42%) were singletons found in only a single genome.

The pangenome is open but approaching closure (Figure 3). Accumulation curves showed each additional genome contributing a mean of 45 new clusters at 84 genomes, with a decay ratio of 0.55. The proportionally large core genome (2.4%) and moderate singleton fraction (42%) are consistent with 9 species spanning a relatively narrow range of genomic diversity.

**Figure 3:**
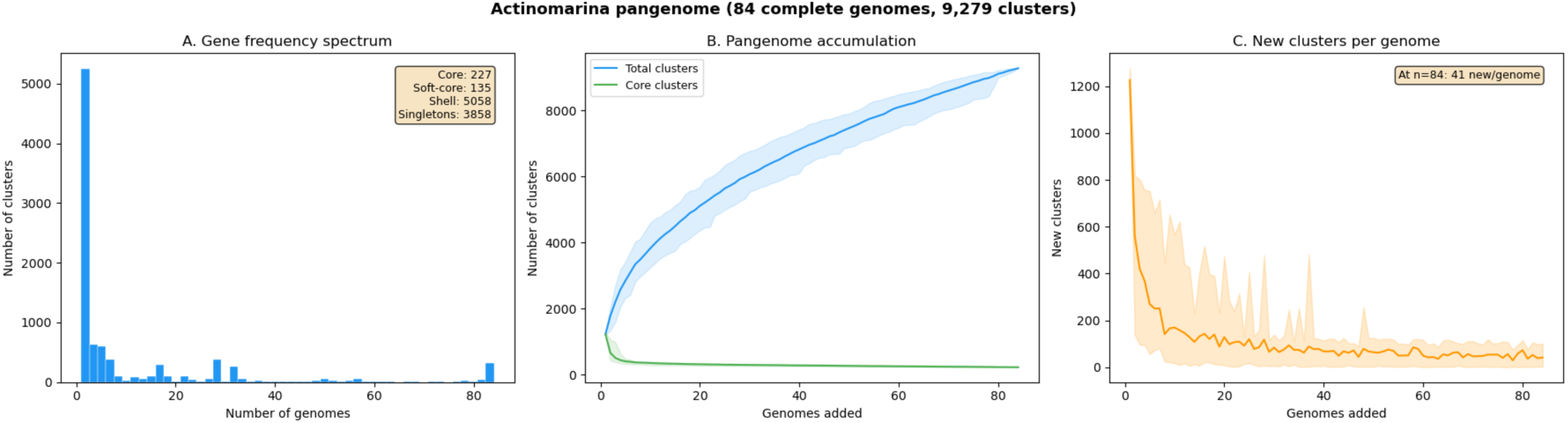
Pangenome overview. (A) Gene frequency spectrum showing core, shell, and singleton categories. (B) Pangenome accumulation curves — total clusters and core decay. (C) New clusters per genome added.

#### Singleton clusters are concentrated in genomic islands

The 3,858 singletons form 724 islands of ≥2 adjacent singleton genes, with 88.4% of all singletons occurring within islands (Supplementary Table 4). Island sizes ranged from 2 to 61 genes (mean 4.7, median 2). Per-genome singleton counts ranged from 1 to 655 (mean 46). Singletons have lower GC content than non-singletons (29.2% vs 32.8%), consistent with recent horizontal acquisition and subsequent amelioration toward the host’s AT-biased composition.

#### A hypervariable region at 85–90% from dnaA

Mapping singleton density along the chromosome (Figure 4) reveals a pronounced hotspot at 85–90% of the genome from dnaA, where 1,019 singleton genes across the 84 genomes are concentrated in a single 5% window — 5.3× the expected density under a uniform distribution (χ² = 5,876, df = 19, p < 10⁻³⁰⁰; 2×2 test for HVR bins vs rest: χ² = 3,830, p < 10⁻³⁰⁰). The 7 largest genomic islands (out of 724; Supplementary Table 5) all map to the 84–90% region. A secondary, weaker hotspot is present at 10–15% (499 singletons).

**Figure 4:**
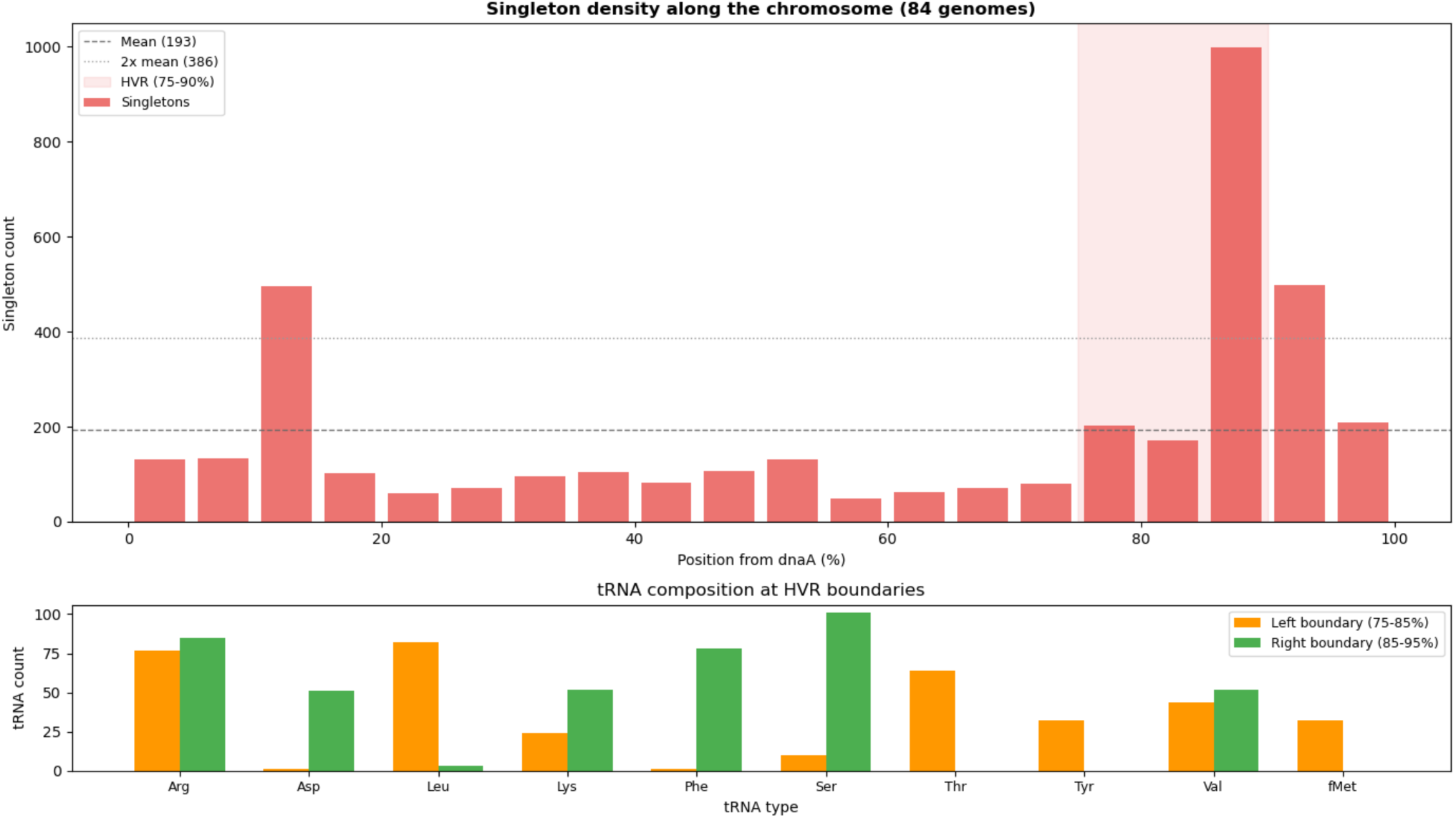
Hypervariable region. Top: singleton density by chromosomal position (5% bins), showing the HVR peak at 85–90% from dnaA with selA position marked. Bottom: tRNA composition at the left and right HVR boundaries.

The primary HVR occupies the same replicative distance from the origin as *Pelagibacter*’s HVR (7–15% from dnaA), but on the opposite replichore: *Pelagibacter*’s HVR is clockwise from dnaA, while *Actinomarina*’s is counterclockwise. Both HVRs are bounded by tRNA genes at both flanks. In *Actinomarina*, the left boundary (79–83% from dnaA) is enriched for Arg, Thr, Tyr, and Met tRNAs, while the right boundary (91–95%) is enriched for Phe, Arg, Lys, and Asp tRNAs — different specific tRNAs from those flanking *Pelagibacter*’s HVR (Phe/His at left, Arg at right), but the same architectural principle of tRNA-bounded integration sites.

The selenocysteine tRNA (selC) is physically located inside the HVR in 32 of 84 genomes. The selenocysteine synthase gene selA maps to the right HVR boundary (91–92% from dnaA) in all genomes. The implications of this co-localization are discussed in the selenocysteine section below.

Lopez-Perez et al. (2020) identified a “flexible genomic island preserved throughout the order” that was “related to the sugar decoration of the envelope” and used “several tRNAs as hot spots to increase its genomic diversity.” Our 84 complete genomes confirm and extend this observation: the island is a positionally conserved HVR at 85–90% from dnaA, bounded by specific tRNA clusters, with 1,019 singleton genes concentrated in a 5% chromosomal window.

Unlike *Pelagibacter*, where the 16S–23S rRNA pair is embedded within the HVR at 8–10% from dnaA, Infernal cmsearch against bacterial 16S, 23S, and 5S covariance models reveals that all three rRNA genes in *Actinomarina* form a compact operon near the replication terminus (44–54% from dnaA), with a 135–153 bp ITS spacer between 16S and 23S and the 5S just 46–49 bp downstream. The rRNA operon is entirely separate from the HVR in all 84 genomes.

#### Assembly frameshifts are rare and mostly biological

We detected 463 frameshifted genes across the 84 genomes (0.5% of all genes, 5.5 per genome; Supplementary Table 6). Of the 334 split genes (adjacent ORFs matching the same UniRef90 target), 48% have no homopolymer run longer than 4 bp at the junction, indicating that the majority of frameshifts are biological (pseudogenes, programmed frameshifts, or overlapping reading frames) rather than basecalling artifacts. The low frameshift rate is consistent with *Actinomarina*’s GC content (32.8%), which produces relatively few long homopolymer runs — the primary source of systematic ONT basecalling errors (Sereika et al., 2022).

### Functional annotation and metabolic capacity

UniRef90 annotation via MMseqs2 identified homologs for 93,577 of 102,662 proteins (91.2%). Actinorhodopsin was detected in all 84 genomes via UniRef90 (best hit: UniRef90_A0A0U2XMT3, actinorhodopsin; 75–78% identity), confirming the photoheterotrophic capacity of the genus. KofamScan assigned KEGG orthologs to 50,747 proteins (693 unique KOs across the genus, mean 604 assignments per genome) at the strict, profile-specific threshold.

### Extreme auxotrophy

Pathway reconstruction from KofamScan assignments (Figure 5; Supplementary Table 7) reveals a pattern of metabolic dependency more extensive than that of *Pelagibacter* (which retains arginine biosynthesis, most tryptophan biosynthesis steps, and thiamine salvage; Lui & Nielsen, in preparation) and, to our knowledge, any other free-living marine bacterium documented from complete genomes.

**Figure 5:**
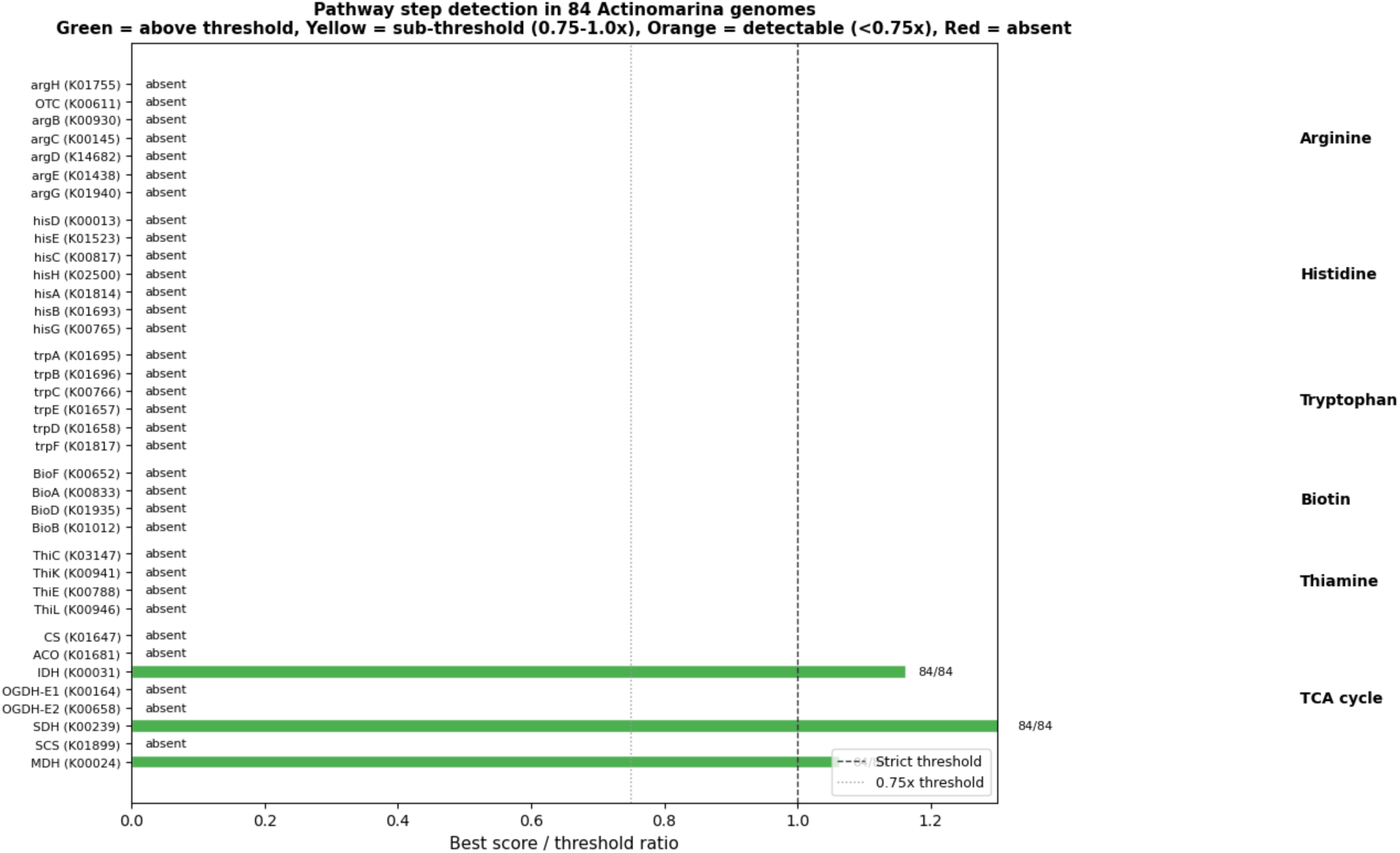
Pathway step detection in 84 Actinomarina genomes. Bars show best KofamScan score as a fraction of profile threshold. Green = above threshold, yellow = sub-threshold (0.75–1.0×), orange = detectable but weak, red = absent.

As a positive control, the same KofamScan pipeline correctly detects arginine (5/7 steps), histidine (6/7), tryptophan (5/6), biotin (4/4), and thiamine (4/4) biosynthesis in *Streptomyces coelicolor* A3(2), confirming that these pathways are detected when present in an Actinobacterium. The missing 1–2 steps in arginine, histidine, and tryptophan reflect alternative enzymes in *S. coelicolor* that fall outside standard KEGG orthology, not KofamScan false negatives.

#### Universally absent pathways (0 of 84 genomes at strict threshold)

Arginine biosynthesis is absent at the strict threshold; one step (K01438, acetylornithine deacetylase) scores at 78% of threshold in 19/84 genomes, but the remaining steps are undetectable. Histidine biosynthesis is absent. Tryptophan biosynthesis is absent. Thiamine biosynthesis is completely absent — not even salvage kinase is retained.

Biotin biosynthesis requires nuanced interpretation. The first committed step, 8-amino-7-oxononanoate synthase (BioF, K00652), scores at 88% of the profile threshold in 24 of 84 genomes — consistent with a divergent homolog, though cross-reactivity to a related aminotransferase family cannot be excluded. However, the final step, biotin synthase (BioB, K01012), scores at only 37% of threshold in any genome, strongly suggesting that the pathway is non-functional regardless of BioF status.

The TCA cycle retains 5 of 8 steps at strict threshold: isocitrate dehydrogenase (K00031, 84/84), succinate dehydrogenase (K00239, 84/84), fumarase (K01679, 83/84), malate dehydrogenase (K00024, 84/84), and succinyl-CoA synthetase alpha subunit (K01902, 84/84). The beta subunit (K01903) scores at 93.5–97.9% of threshold in all 84 genomes — almost certainly present but technically sub-threshold. The dihydrolipoyl dehydrogenase E3 subunit (K00382) also passes strict threshold in 83/84 genomes, though this subunit is shared with pyruvate dehydrogenase and does not on its own indicate OGDH function. The dihydrolipoamide succinyltransferase E2 component of the OGDH complex (K00658) has a divergent homolog below threshold (75–80% of threshold in 78/84 genomes). Citrate synthase, aconitase, and 2-oxoglutarate dehydrogenase E1 are genuinely absent (best scores 2–67% of threshold). The cycle is therefore incomplete at the entry point (no citrate synthase or aconitase) but retains most of the oxidative branch from isocitrate through to oxaloacetate. The universal retention of isocitrate dehydrogenase and isocitrate lyase in the absence of the enzymes that produce isocitrate raises the question of how these enzymes obtain their substrate. Whether an undetected aconitase variant, environmental uptake, or an alternative metabolic route supplies isocitrate remains to be determined.

#### Universally retained pathways

GlyA (serine hydroxymethyltransferase, K00600) is present in all 84 genomes. One of the two glyoxylate bypass enzymes, isocitrate lyase, is present in all 84 genomes. Serine biosynthesis is partially retained (serA and serB present, serC absent).

#### Variable pathways

The diaminopimelate pathway for lysine biosynthesis (K00215) is present in only 40 of 84 genomes. Glyoxylate cycle malate synthase (K01638) is present in 20 of 84. Assimilatory sulfate reduction (K00957) is present in only 6 of 84, with the remaining steps absent.

*Actinomarina*’s auxotrophies are largely uniform across the genus. The four universally absent pathways (arginine, histidine, tryptophan, thiamine) are absent in all 9 species without exception, and biotin is non-functional in all. This uniformity suggests that these losses occurred in the common ancestor of the genus and have been fixed ever since, rather than representing ongoing, lineage-specific pathway erosion.

Lopez-Perez et al. (2020) suspected extreme auxotrophy in *Actinomarina* on the basis of SAG data but could not confirm it, because every apparent gene absence in a SAG or MAG might reflect incomplete genome recovery. Our 84 complete genomes make every absence definitive: the genes are not present in any copy, in any species, at any genomic position.

*Actinomarina* survives via rhodopsin-based photoheterotrophy — light-driven proton pumping supplements energy from organic carbon oxidation — supplemented by scavenging dissolved amino acids, vitamins, and other metabolites from the environment. Consistent with this dependence on exogenous nutrients, most genomes encode ABC transporters for branched-chain amino acids (82/84), polar amino acids (74/84), and biotin (K03523, 84/84), providing import capacity for the nutrients they cannot synthesize.

To complement sequence-based annotation, we predicted structures for 9,221 of the 9,278 cluster representative proteins using ESMFold (57 exceeded 1,035 residues, the memory capacity of our GPU) and searched them against the AlphaFold Database using Foldseek. Of these, 7,653 (83.0%) matched an AlphaFold entry (E-value ≤ 10⁻³). Cross-referencing sequence and structural annotation across the 9,278 cluster representatives reveals their complementary value: 6,452 (69.5%) were identified by both methods, 314 (3.4%) by sequence only, and 1,201 (12.9%) by structure only — functional information that sequence similarity alone could not provide. A total of 1,311 proteins (14.1%) had no detectable match by either method. Of these, 92 are conserved in 10 or more genomes — proteins present across most species that match nothing in any database and may warrant experimental characterization.

### Gene order is not conserved across species

Gene order conservation was assessed by pairwise adjacency analysis across all 84 complete genomes (Figure 6). Of 13,324 unique gene adjacencies (defined as consecutive genes in the same orientation on the same strand), zero are universal — not a single adjacency is conserved in all 84 genomes. Only 5 adjacencies are near-universal (present in ≥80 of 84 genomes), and 42% of all adjacencies (5,594) occur in only a single genome.

**Figure 6:**
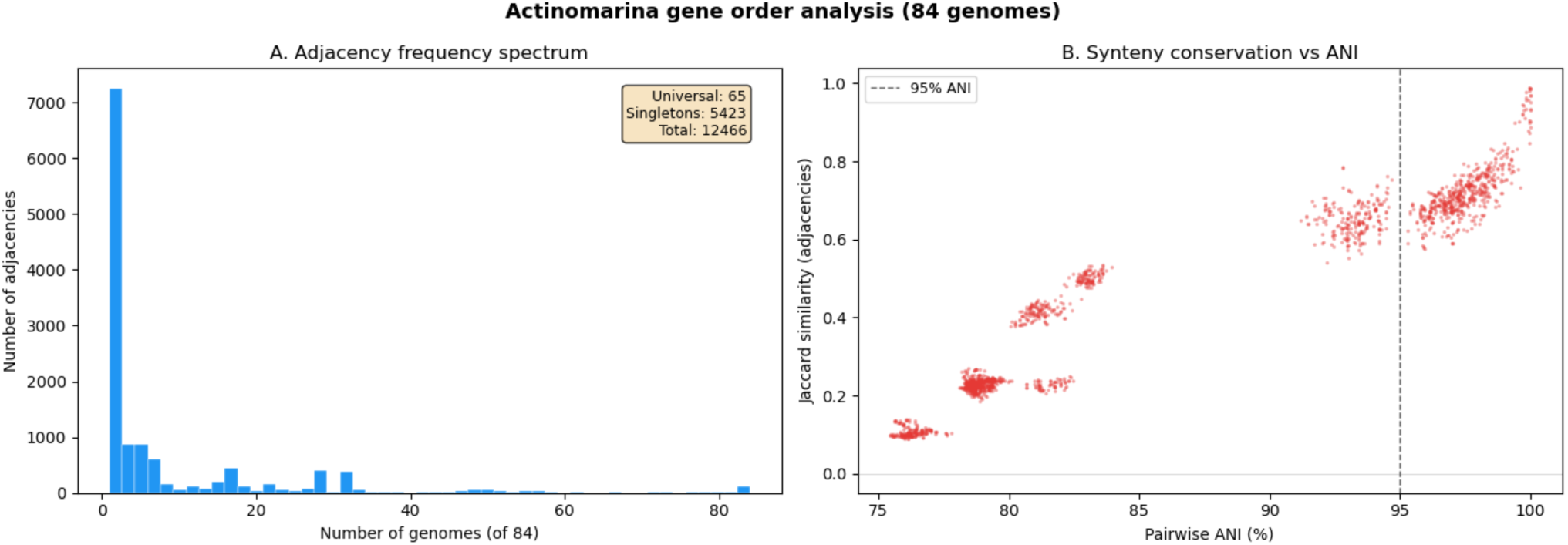
Gene order analysis. (A) Adjacency frequency spectrum — zero universal adjacencies across 84 genomes. (B) Pairwise synteny Jaccard similarity vs ANI — sharp decay of gene order conservation below 95% ANI.

Within-species gene order conservation, measured as pairwise Jaccard similarity of adjacency sets (Fan et al., 2025), is moderate (0.73), while between-species conservation drops sharply (0.24). The pattern is consistent with operonic blocks that are conserved within species but shuffled between species over longer evolutionary timescales. The rearrangement rate is fast enough to disrupt essentially every gene pair given sufficient evolutionary divergence between species.

The scarcity of intrachromosomal repeats (most genomes have only 2, maximum length 777 bp) makes homologous recombination between internal repeats an unlikely rearrangement mechanism. Other possibilities include illegitimate (non-homologous) recombination, recombination at tRNA sites, and phage-mediated rearrangements — the same tRNA-targeted integration that builds the HVR could, over longer timescales, shuffle adjacent gene blocks.

### Universal selenocysteine machinery

All 84 complete *Actinomarina* genomes encode selenocysteine tRNA (selC) with the characteristic UCA anticodon. This finding was confirmed by tRNAscan-SE in bacterial mode across all genomes without exception. In addition to selC, KofamScan identifies selenocysteine synthase (selA) and selenophosphate synthetase (selD) in all 84 genomes (Supplementary Table 8). The selenocysteine-specific translation elongation factor selB is present in all 84 genomes but scores below the KofamScan profile threshold (best score 268, threshold 306), consistent with sequence divergence from the profile training set rather than genuine absence.

Because selD serves both the selenocysteine (Sec) and selenouridine (SeU) pathways, the presence of selD alone would be ambiguous. However, the SeU pathway requires the tRNA 2-selenouridine synthase YbbB (K06917), and Peng et al. (2016) showed that the SeU trait is absent in all sequenced Actinobacteria. We confirm this: K06917 is not detected in any of the 84 genomes above the profile threshold (best score 33.9 vs threshold 127.6), and no selenouridine-related proteins appear in UniRef90 or Foldseek results. The co-occurrence of selA, selC, and selD with the absence of YbbB establishes that the Sec machinery in *Actinomarina* is for selenocysteine incorporation, not selenouridine modification.

This finding has not been previously reported. Lopez-Perez et al. (2020), the most comprehensive genomic study of the Actinomarinales to date, did not mention selenocysteine or any component of the Sec machinery. Surveys of selenoprotein utilization across bacterial phyla (Zhang et al., 2006; Zhang & Gladyshev, 2008) have shown that Sec usage is widespread but patchy. Universal selenocysteine usage across all 9 species and all 84 genomes in a genome of ∼1.1 Mbp is, to our knowledge, not previously documented in any abundant marine picoplankton lineage.

Selenocysteine (Sec) is the 21st genetically encoded amino acid, incorporated at UGA codons that would otherwise signal translation termination. Sec residues are catalytically superior to cysteine in redox enzymes, with selenoenzymes that can be over 100× more catalytically efficient than their cysteine homologs (Arnér, 2010; Kim & Gladyshev, 2005). The retention of dedicated Sec biosynthetic machinery (selA, selB, selC, selD) in a genome of ∼1.1 Mbp implies that the catalytic advantage is large enough to justify the cost. In a photoheterotrophic organism that inhabits the sunlit ocean surface, where reactive oxygen species (ROS) are generated continuously by UV radiation and photochemistry, deep-Sep prediction identifies ∼5 selenoproteins per genome, including a selenoprotein form of selD itself. The majority of the selenoprotein families are uncharacterized, and their specific functions remain to be determined experimentally.

The chromosomal context of the Sec machinery is unusual (Figure 7). selC maps to the HVR (85–90% from dnaA) in 32 of 84 genomes, while it occupies a position outside the HVR in the remaining 52. selA maps consistently to the right HVR boundary (91–92% from dnaA) in all genomes. The variable position of selC — sometimes inside, sometimes outside the HVR — suggests that selC may be subject to the same tRNA-site-targeted integration and recombination that drives HVR content turnover. This would be consistent with the known role of tRNA genes as integration hotspots for phage and genomic islands.

**Figure 7:**
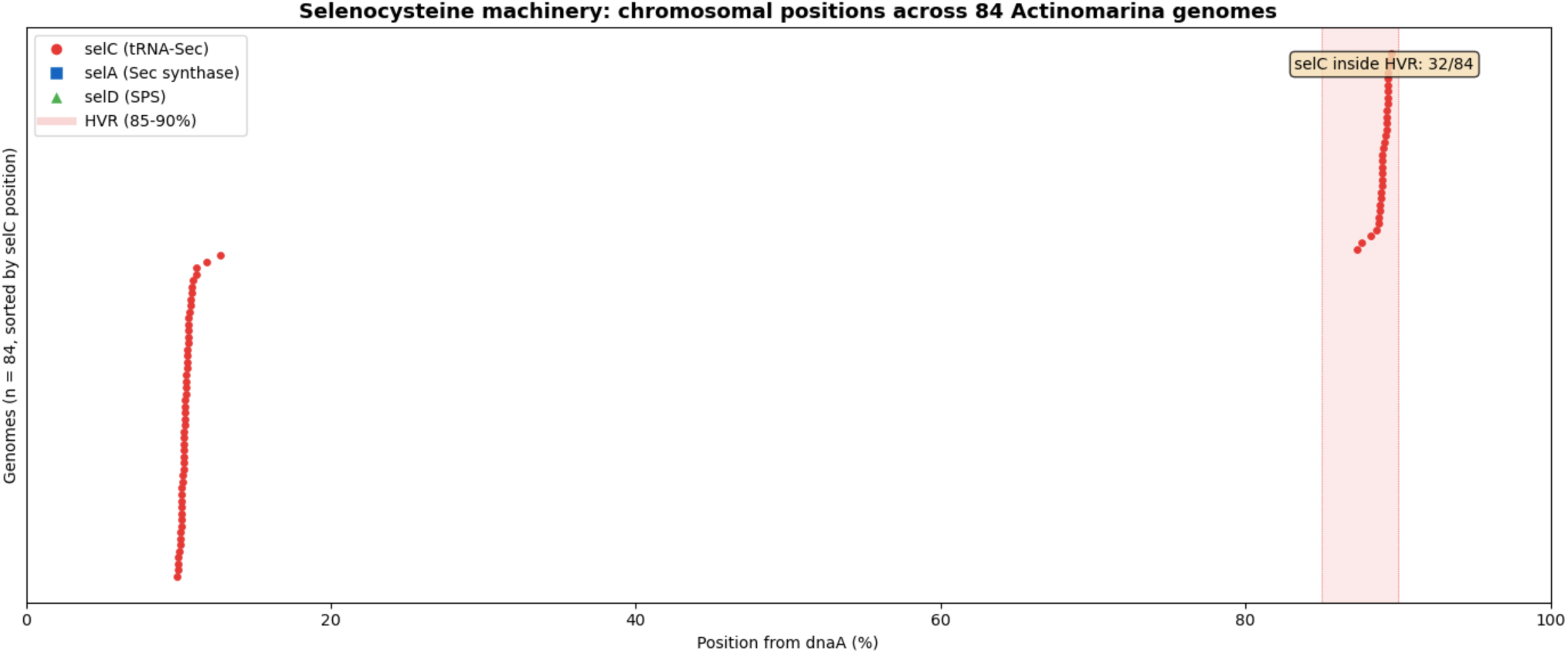
Selenocysteine machinery — chromosomal positions of selC (tRNA-Sec), selA (Sec synthase), and selD (selenophosphate synthetase) across 84 genomes, with HVR region shaded. Genomes sorted by selC position.

We searched all 113 genomes with four Rfam SECIS covariance models (RF00031, RF01988, RF01989, RF01990) using Infernal cmsearch. Only 6 hits were obtained, all with E-values above the significance threshold (6.1–81). The best hits (E = 6.1) were SECIS_1 model matches inside characterized genes in two genomes, in the same genomic context flanked by ATP synthase genes. Two additional hits mapped to the HVR region (86.7% and 94.6% from dnaA). The failure of these models is expected: RF00031 is primarily a eukaryotic SECIS model, and bacterial selenocysteine insertion sequences (bSECIS) are structurally distinct stem-loops immediately downstream of the UGA codon, which are poorly represented in Rfam.

To identify the selenoprotein targets directly, we applied deep-Sep, a deep learning tool for bacterial selenoprotein gene prediction, to all 113 genomes. deep-Sep identified 550 selenoprotein candidates across all 113 genomes (mean 4.9 per genome, range 3–6), each containing a single selenocysteine (U) residue. These group into 69 protein families based on N-terminal sequence similarity (first 20 residues). The most abundant family (59 genomes, 78 aa; with a closely related variant in 35 additional genomes) is uncharacterized — it has no UniRef90 hit and no KofamScan assignment. The selD family (MRLT* prefix, 342–346 aa, 56 members across 4 sequence variants) maps to selenophosphate synthetase (K01008), indicating that *Actinomarina*’s selD exists in a selenoprotein form in approximately half of genomes, with the remainder presumably carrying the cysteine variant — a known polymorphism first identified by Guimarães et al. (1996). A third major grouping (49 genomes) matches K07090, a conserved protein of unknown function. Of the 550 candidates, 83 (15%) did not overlap any Pyrodigal-predicted gene on the same strand — these fall in regions where standard gene callers treat UGA as a stop codon rather than a selenocysteine codon. As a negative control, applying deep-Sep to 135 complete *Pelagibacter* genomes, which lack selC, returned zero selenoprotein candidates. The identification of ∼5 selenoproteins per genome in *Actinomarina* confirms the functional significance of the Sec biosynthetic machinery and explains the selective pressure to maintain it. Applying deep-Sep to the 23 NCBI *Actinomarina* genomes identified 116 additional candidates (mean 5.0 per genome), with 24 of 69 protein families shared between SFE and NCBI genomes, confirming that selenoprotein usage is a genus-wide feature and not specific to the SFE.

Marine surface waters provide sufficient selenium to support selenoprotein usage. Dissolved selenium in the ocean ranges from 0.5 to 3 nmol/L, with organic selenide comprising up to 80% of total dissolved Se at productivity maxima (Cutter & Bruland, 1984). The San Francisco Estuary has measured dissolved Se concentrations of ∼0.91 nmol/L (Cutter & Cutter, 2004). Selenium availability is therefore not a limiting factor for selenoprotein usage in an environment where *Actinomarina* is abundant.

## Discussion

### Complete genomes for an entire order

The 84 complete genomes presented here are the first for any member of the order Actinomarinales, which is represented in NCBI exclusively by fragmented MAGs. Our genomes outnumber the confirmed high-quality MAGs at NCBI by nearly 4:1, and the 41% misclassification rate among NCBI genomes that pass quality thresholds underscores the danger of relying on uncurated public databases for taxonomic or functional inference.

Complete genomes enabled three discoveries that were invisible to prior SAG- and MAG-based studies: the universal selenocysteine machinery, the tRNA-bounded hypervariable region, and the definitive confirmation of auxotrophies that Lopez-Perez et al. (2020) could only hypothesize. Each of these depends on the ability to confirm presence or absence of specific genes across the entire chromosome, which is only possible with complete genomes.

### Selenocysteine: catalytic advantage in a small-genome photoheterotroph

The retention of selenocysteine capacity in all 84 genomes, across all 9 species, implies that the selective advantage is large and universal within the genus. In a genome of ∼1.1 Mbp, a dedicated biosynthetic pathway (selA, selB, selC, selD) is a substantial investment. The most parsimonious explanation is that one or more selenoproteins provide an advantage that outweighs the cost of maintaining the Sec pathway.

The deep-Sep analysis identifies ∼5 selenoproteins per genome in 69 families, confirming that this investment serves multiple targets. Notably, selD itself is a selenoprotein in *Actinomarina* — the active-site cysteine is replaced by selenocysteine, a variant first identified by Guimarães et al. (1996) in which selenocysteine replaces cysteine at the enzyme’s active site. The majority of the selenoprotein families, however, are uncharacterized and have no functional annotation in any database. Their identification as selenoproteins is a first step; determining their biochemical roles will require experimental work.

The photoheterotrophic lifestyle of *Actinomarina* provides a plausible context for why selenoproteins are advantageous. Rhodopsin-mediated proton pumping occurs at the cell surface in the photic zone, where UV radiation and photochemically generated ROS are constant threats. Selenocysteine-containing redox enzymes can be over 100× more efficient than their cysteine analogs at neutralizing ROS (Arnér, 2010). If even one of the ∼5 selenoproteins per genome is a ROS-defense enzyme, the fitness benefit could exceed the cost of maintaining the Sec pathway.

### Convergent genome architecture, divergent metabolic solutions

*Actinomarina* and *Pelagibacter* are two abundant marine bacterial lineages from different phyla (Actinomycetota and Pseudomonadota). Despite this phylogenetic distance, they share remarkably similar genome architectures: small genomes (1.1 vs 1.3 Mbp), few internal repeats, high coding density (>95%), and a tRNA-bounded hypervariable region at a similar replicative distance from the origin (7–15% from dnaA in *Pelagibacter*, 85–90% — i.e. 10–15% counterclockwise — in *Actinomarina*). The HVR in both genera carries genome-specific content — likely surface modification genes involved in phage evasion — flanked by tRNA integration sites. Both HVRs also show a coding-strand bias toward the leading strand of their respective replichore: 68% plus-strand in *Pelagibacter*’s HVR (right replichore, 7–15% from dnaA) and 61% minus-strand in *Actinomarina*’s (left replichore, 85–90%), compared to ∼50% in the rest of each genome. This shared bias likely reflects selection against head-on collisions between the replication and transcription machineries.

The rRNA genes, however, occupy entirely different contexts. In *Pelagibacter*, the 16S–23S pair is embedded within the HVR and the 5S is separated by ∼47 kbp; in *Actinomarina*, all three rRNA genes form a compact operon (16S–ITS–23S–5S within ∼200 bp) at the replication terminus, far from the HVR. The convergence extends to HVR architecture but not to rRNA organization.

Yet the metabolic strategies are substantially different. *Pelagibacter* retains arginine biosynthesis and most tryptophan biosynthesis steps and varies in its auxotrophic profile across species. *Actinomarina* retains an incomplete TCA cycle (5 of 8 steps at strict threshold, but lacking the entry enzymes citrate synthase and aconitase), cannot synthesize arginine, histidine, or tryptophan, and shows no species-level variation in these dependencies. Where *Pelagibacter*’s auxotrophies are phylogenetically structured — still evolving — *Actinomarina*’s appear to be fixed in a minimal metabolic configuration that is supplemented by photoheterotrophic energy acquisition and dependent on scavenging organic nutrients.

The pangenome reflects this difference: *Actinomarina*’s is approaching closure (decay ratio 0.55) while *Pelagibacter*’s remains wide open (0.94; Lui & Nielsen, in preparation). Fewer species, less HGT, and a larger proportional core suggest that *Actinomarina* occupies a narrower ecological niche — or that a genome this small leaves less room for accessory gene content to vary.

### The HVR as a convergent feature of small-genome marine bacteria

The discovery of tRNA-bounded HVRs at similar replicative distances from the origin in both *Actinomarina* (85–90% from dnaA, i.e. ∼10–15% counterclockwise) and *Pelagibacter* (7–15% clockwise) suggests that this architecture may be a general feature of small-genome marine bacteria. In companion studies (Lui & Nielsen, in preparation), we report tRNA-bounded HVRs in two additional genera — *Planktophila* and D2472/SAR86 — at different chromosomal positions. The tRNA-bounded architecture is shared across all four genera; whether the replicative distance is also conserved will require further analysis.

The mechanistic basis for the tRNA-boundary architecture is likely the well-documented role of tRNA genes as integration hotspots for phage and genomic islands. Why HVRs cluster within the first quarter of a replichore in *Actinomarina* and *Pelagibacter* remains unknown. Testing this will require identifying the integration mechanism and the selective forces that maintain HVR content.

## Methods

### Sample collection and sequencing

Water samples were collected from 8 stations along the SFE salinity gradient in summer (July–August 2022) and winter (January–March 2023). Each sample was sequenced on two PromethION flow cells (ONT R10.4 chemistry). Detailed sampling and sequencing protocols are described in Lui & Nielsen (2024).

### Metagenome assembly

Metagenome assembly was performed with myloasm v0.4.0 (Shaw, Marin & Li, 2026) using default parameters for ONT R10 data. Circular contigs were identified by myloasm’s circularization detection.

### *Actinomarina* genome identification and quality assessment

All contigs ≥500 kbp were assessed with CheckM2 v1.1.0 (Chklovski et al., 2023). High-quality genomes were defined as ≥90% estimated completeness and <5% estimated contamination. Taxonomic classification was performed with GTDB-Tk v2.6.1 (Chaumeil et al., 2022) using the GTDB R226 reference database (Parks et al., 2025). Genomes classified as g Actinomarina were retained.

### NCBI reference genomes

All 396 *Actinomarina* genome assemblies were downloaded from NCBI (March 2026) using the NCBI Datasets CLI v18.10.2. Quality was assessed with CheckM2; 39 passed high-quality thresholds. GTDB-Tk reclassification identified 23 as genuinely *Actinomarina* (16 misclassified to other genera). Pairwise ANI to our 84 genomes was computed with skani.

### Genome reorientation

Complete genomes were reoriented to begin at the dnaA gene using dnaapler v1.3.0 (Bouras et al., 2024) in bulk chromosome mode. Non-circular genomes were padded with 10 N’s at contig boundaries, pseudo-circularized, and reoriented; 27 of 29 succeeded, and 2 failed due to absent dnaA.

### Gene prediction

Protein-coding genes were predicted with Pyrodigal v3.6.3 (Larralde, 2022) in single-genome mode (-p single). Single-genome mode was used rather than metagenomic mode to train gene-calling parameters on each genome’s specific codon usage and start-site patterns.

### tRNA prediction

tRNA genes were predicted with tRNAscan-SE v2.0.12 (Chan & Lowe, 2019) in bacterial mode (-B). Selenocysteine tRNAs were identified by the diagnostic UCA anticodon.

### rRNA gene identification

Ribosomal RNA genes were identified using Infernal v1.1.5 (Nawrocki & Eddy, 2013) cmsearch against three Rfam covariance models: RF00177 (bacterial SSU/16S), RF02541 (bacterial LSU/ 23S), and RF00001 (5S). All 84 rotated genomes were searched with 32 CPUs and –noali. Chromosomal positions were expressed as fractions of genome length from dnaA. The ITS spacer between 16S and 23S and the distance from the 23S to 5S were computed from hit coordinates to assess operon structure.

### Average nucleotide identity

Pairwise ANI values were computed with skani v0.3.1 (Shaw & Yu, 2023) in triangle mode using the –slow preset. Species-level clusters were defined at 95% ANI using single-linkage connected components.

### Phylogenomic analysis

Single-copy core protein clusters (present in exactly one copy in all 84 complete genomes) were identified from the pangenome clustering. Each cluster was aligned with MAFFT v7.526 (Katoh & Standley, 2013) using –auto. Aligned sequences were concatenated into a supermatrix of 65,482 amino acid positions across 227 partitions. Phylogenomic inference was performed with IQ-TREE v3.0.1 (Minh et al., 2020; Wong et al., 2025) with per-partition substitution models selected by ModelFinder (Kalyaanamoorthy et al., 2017) — LG+F+G4 was selected for all 227 partitions — and 1000 ultrafast bootstrap replicates (Hoang et al., 2018).

For the expanded tree, protein sequences from 29 non-circular genomes and 23 NCBI genomes were searched against the 227 core cluster representatives using MMseqs2 (minimum 30% identity, E-value ≤ 10⁻⁵, 50% coverage). Best-hit sequences were added to existing alignments using MAFFT –add –keeplength. Genomes lacking a core gene received an all-gap sequence for that partition.

The tree was rooted using 3S_u11376018 (Casp-actino5, order Acidimicrobiales), which retains 164 of 227 core genes at ≥30% amino acid identity and 50% coverage.

### Pangenome analysis

Predicted proteins from all 84 complete genomes were clustered with MMseqs2 v18.8cc5c (Steinegger & Söding, 2017) easy-cluster at 70% sequence identity and 80% bidirectional coverage. Pangenome accumulation curves were computed from 50 random genome addition orders. The decay ratio was calculated as the ratio of new clusters added by the last genome to new clusters added by the second genome.

### Assembly frameshift detection

Split genes were detected as adjacent ORFs in the same genome matching the same UniRef90 target with an intergenic distance <50 bp. Truncated-with-gap frameshifts were detected as proteins less than 80% of their UniRef90 match length with a downstream intergenic gap exceeding 50 bp. Homopolymer runs at each junction were measured, and frameshifts were classified as inherited biological (same gene split in multiple genomes of the same species), convergent biological (same gene split in unrelated species), or technical (unique to a single genome).

### Functional annotation

Proteins were searched against UniRef90 (downloaded March 2026) using MMseqs2 easy-search with minimum 30% identity, E-value ≤10⁻⁵, and 50% coverage. KEGG ortholog assignments were made using KofamScan v1.3.0 (Aramaki et al., 2020) with KEGG HMM profiles (downloaded March 2026) and profile-specific adaptive thresholds. The strict, profile-specific threshold was used for primary pathway assignments; sub-threshold hits are reported individually where relevant (Supplementary Table 7). As a positive control, the same pipeline was applied to the *Streptomyces coelicolor* A3(2) proteome (GCF_000203835.1; 7,769 proteins), a model Actinobacterium with experimentally characterized biosynthetic pathways.

### Structural annotation

Protein structures were predicted for 9,221 cluster representative proteins using ESMFold (Lin et al., 2023) on an NVIDIA RTX 6000 Ada Generation GPU. Predicted structures were searched against the AlphaFold Database v4 (Varadi et al., 2024) using Foldseek v10.941 (van Kempen et al., 2024) with E-value ≤ 10⁻³, sensitivity 9.5, structural alignment mode (–alignment-type 2), and up to 5 hits per query.

### Singleton and island characterization

Singleton clusters (present in exactly one genome) were characterized by cross-referencing against UniRef90 annotations. Genomic islands were detected as maximal runs of ≥2 consecutive singleton genes. Per-gene GC content was calculated from nucleotide sequences.

### Hypervariable region analysis

Singleton density was computed in 5% bins along the chromosome (fraction of genome length from dnaA). The HVR was defined as the contiguous set of bins with singleton density exceeding 2× the genome-wide mean. tRNA genes at HVR boundaries were identified from tRNAscan-SE output. Positions of selA, selC, and selD relative to dnaA were extracted from GFF annotations and KofamScan assignments. Non-random singleton clustering was tested with a chi-squared goodness-of-fit test against the null hypothesis of uniform distribution across 20 bins (expected count per bin = total singletons / 20). A 2×2 chi-squared test of independence was used to compare singleton frequency in the HVR bins versus all other bins.

### Selenocysteine machinery identification

Selenocysteine tRNA (selC) was detected by tRNAscan-SE. Selenocysteine synthase (selA, K01042) and selenophosphate synthetase (selD, K01008) were identified by KofamScan at the strict adaptive threshold. The selenocysteine-specific elongation factor selB (K03833) was detected in all 84 genomes below the KofamScan adaptive threshold (best score 268, threshold 306, 87.6%); its consistent presence across all genomes at this level, combined with the functional requirement for selB in Sec incorporation, supports its identification as a divergent but genuine homolog. Selenoprotein gene prediction was performed with deep-Sep v1.0 (Xiao & Zhang, 2025), a transformer-based deep learning tool for bacterial selenoprotein ORF identification. All 113 SFE genome FASTAs, the 23 NCBI *Actinomarina* genome FASTAs, and 135 *Pelagibacter* genomes (as a negative control lacking selC) were submitted as separate runs. deep-Sep identifies ORFs containing in-frame UGA codons and classifies them as selenocysteine-encoding based on sequence context. Predicted selenoprotein ORFs were mapped to Pyrodigal gene predictions by genomic coordinate overlap (same strand, maximum overlap). Selenoprotein families were defined by grouping predictions with identical first 20 amino acid residues.

SECIS elements were searched using Infernal v1.1.5 (Nawrocki & Eddy, 2013) cmsearch with Rfam 14.10 (Kalvari et al., 2021) covariance models RF00031 (SECIS_1), RF01988 (SECIS_2), RF01989 (SECIS_3), and RF01990 (SECIS_4), with default E-value threshold (10.0).

### Genome synteny

Gene adjacencies were defined as consecutive genes sharing the same orientation, represented as pairs of cluster IDs with relative strand information. Adjacencies were canonicalized (smaller cluster ID first). Pairwise Jaccard similarity was computed as | intersection|/|union| of adjacency sets. Frequency spectra and within- vs between-species comparisons used the 84 complete genomes.

### Repeat analysis

Intra-genomic repeats were identified by nucmer v3.23 (Kurtz et al., 2004) self-alignment (each genome against itself) with default parameters (minimum match length 20 bp). Maximum repeat length and repeat count per genome were recorded.

## Data availability

All genome sequences and annotations have been deposited at [repository TBD]. Assembly and annotation scripts are available at [repository TBD].

## Acknowledgments

We thank Erica Nejad and the crew of USGS R/V David H. Peterson for collecting samples. We cannot overemphasize the value of the support we have received from them. We also thank Miten Jain for lively discussions about nanopore sequencing and providing support for the PromethION sequencing.

Torben also wants to thank Lauren for putting up with how he does things; it can be challenging.

## Supplementary Information

**Supplementary Table 1:** Genome statistics for all 113 *Actinomarina* genomes — Genome ID, Station, Season, Circular (Y/N), Genome size (bp), GC (%), Gene count, Coding density (%), Completeness (%), Contamination (%), GTDB-Tk classification, Species cluster (1–9), tRNA count, selC present (Y/N).

**Supplementary Table 2:** NCBI *Actinomarina* comparison — 396 genomes assessed: accession, assembly level, CheckM2 quality, GTDB-Tk classification, genuine *Actinomarina* (Y/N), best-hit ANI to our genomes, species match.

**Supplementary Table 3:** Core gene list — 227 single-copy core clusters with UniRef90 annotations, KO assignments, alignment length, and mean identity.

**Supplementary Table 4:** Singleton characterization — functional categories, island membership, GC content, and per-genome counts.

**Supplementary Table 5:** HVR characterization — left and right boundary tRNAs, HVR size (genes and bp), singleton count, for all 84 genomes.

**Supplementary Table 6:** Frameshift analysis — per-genome split gene counts, truncated-with-gap counts, homopolymer association, recurrence class.

**Supplementary Table 7:** KofamScan pathway completeness — KEGG module completeness per genome for 84 complete genomes.

**Supplementary Table 8:** Selenocysteine machinery positions — selA, selC, selD positions as fraction of genome from dnaA, selC inside HVR (Y/N), for all 84 genomes.

## Notes

### Competing Interest Statement

The authors have declared no competing interest.

